# Eyes are essential for magnetoreception in a mammal

**DOI:** 10.1101/2020.05.07.082388

**Authors:** Kai R. Caspar, Katrin Moldenhauer, Regina E. Moritz, E. Pascal Malkemper, Sabine Begall

**Affiliations:** Department of General Zoology, University of Duisburg-Essen, Universitaetsstr. 5, 45117 Essen, Germany; Max Planck Research Group Neurobiology of Magnetoreception, Center of Advanced European Studies and Research (CAESAR), Ludwig-Erhard-Allee 2, 53175 Bonn, Germany; Department Vision, Visual Impairment & Blindness, Faculty 13, Technical University of Dortmund, Emil-Figge-Straße 50, 44227 Dortmund, Germany; Department of Game Management and Wildlife Biology, Faculty of Forestry and Wood Sciences, Czech University of Life Sciences, 16521 Praha 6, Czech Republic

**Keywords:** magnetic sense, mole-rat, Bathyergidae, sensory biology, magnetite, animal navigation

## Abstract

Several groups of mammals use the Earth’s magnetic field for orientation, but their magnetosensory organ remains unknown. The Ansell’s mole-rat (*Fukomys anselli*) is a subterranean rodent with innate magnetic orientation behavior. Previous studies proposed that its magnetoreceptors are located in the eye. To test this hypothesis, we assessed magnetic orientation in enucleated mole-rats.

Initially, we demonstrate that enucleation of mole-rats does not lead to changes in routine behaviors. We then studied magnetic compass orientation by employing a well-established nest building assay. To ensure that directional responses were based on magnetic parameters, we tested animals under four magnetic field alignments. In line with previous studies, control animals exhibited a significant preference to build nests in magnetic south-east. In contrast, enucleated mole-rats built nests in random magnetic orientations, suggesting an impairment of their magnetic sense. The results provide robust support for the hypothesis that mole-rats perceive magnetic fields with their minute eyes.

## Introduction

Magnetoreception, the ability to perceive magnetic fields, is a sensory modality that occurs in all major vertebrate groups (Wiltschko & Wiltschko, 1995). The hunt for a magnetic sense in mammals has gained pace since the late 1980s. By now magnetoreception has been shown in numerous taxa, including rodents (Burda et al., 1990; Phillips et al., 2013; Malkemper et al., 2015) and bats (Holland et al., 2006). Even though rodents are readily available for experiments under controlled laboratory conditions, only a few attempts have been made to explore the underlying physiology of this sensory modality in mammals, for example by means of electrophysiology (Semm et al., 1980), pharmacological inhibition (Wegner et al., 2006), or by neural activity mapping (Němec et al., 2001; Burger et al., 2010), contrasting with a great number of such approaches in birds (cf. Mouritsen & Hore, 2012; Wiltschko & Wiltschko, 2013). As a consequence, the location, structure and functional properties of mammalian magnetoreceptors are still elusive (Begall et al., 2014).

Two main mutually non-exclusive magnetoreception mechanisms have been postulated for mammals and other terrestrial vertebrates (Freake & Phillips, 2005; Mouritsen & Hore, 2012). The hypothesis of light-dependent magnetoreception based on radical pairs proposes that photochemical reactions, most likely mediated by retinal blue light-sensitive proteins of the cryptochrome family, are altered by alignment to a magnetic field. Experimental evidence supports a radical-pair mechanism in murine rodents (Phillips et al., 2013; Malkemper et al., 2015; Painter et al., 2018), but whether mammalian cryptochromes are involved is unclear, since they apparently lack the ability to respond to light (Kutta et al., 2017). An alternative hypothesis proposes that magnetite crystals (Fe_3_O_4_) linked to mechanosensitive ion channels lead to magnetically modulated neuronal excitation (Kirschvink & Gould, 1981). In contrast to photochemical transduction systems, magnetite receptors are light-independent and could be located anywhere in the body. In mammals, there is evidence consistent with magnetite-mediated magnetoreception in bats (Wang et al., 2007; Holland et al., 2008) and in Ansell’s mole-rat (*Fukomys anselli*), a subterranean rodent.

Ansell’s mole-rats are a model species in the study of magnetoreception because of their strong innate preference to build nests in the magnetic south-eastern sector of a circular arena (Burda et al., 1990; Marhold et al., 1997a; Thalau et al., 2006; Wegner et al., 2006). Nest building assays have revealed several properties of the mole-rat magnetic compass: it works in total darkness, responds to changes in field polarity but not in inclination, is affected by strong magnetic pulses (Marhold et al., 1997a, b), but is insensitive to radiofrequency magnetic fields (Thalau et al., 2006). These characteristics fit magnetite-based receptors, but their anatomical locus remains unknown.

The eye has a crucial role in the avian magnetic compass response (Mouritsen & Hore, 2012) and has therefore attracted attention as a potential location of magnetoreceptors in mammals (e.g., Wegner et al., 2006; Nießner et al., 2016). Although Ansell’s mole-rat spends most of its life in darkness, its minute eyes display all features typical for epigeic mammals (Peichl et al., 2004; Nemec et al., 2008). Wegner et al. (2006) reported that local lidocaine anesthesia of the cornea led to a loss of directional nest building in Ansell’s mole-rats. The treatment did not impair light-dark discrimination, suggesting that the retina was unaffected by corneal lidocaine application (Wegner et al., 2006). The authors concluded that the mole-rats’ magnetoreceptors are located in the cornea, but the utility of lidocaine for behavioral testing has significant drawbacks. First, the induced anesthesia, even after repeated applications, only lasts up to 15 min (Schønemann et al., 1992), which limits its validity in prolonged behavioral assays (Wegner et al. (2006) allowed nest building for 30-60 min). Additionally, lidocaine is lipophilic and can diffuse widely, potentially affecting untargeted tissues (Engels et al., 2018). Lidocaine further binds to serum proteins in the blood and rapidly crosses the blood-brain barrier, where it may induce non-specific effects on the central nervous system (Pardridge et al., 1983). Therefore, ablation experiments should be preferred over local anesthesia in experiments aimed to narrow down the site of magnetoreceptors (Engels et al., 2018). To test the hypothesis that the magnetoreceptors of Ansell’s mole-rats are indeed located in the eye, we assessed magnetic orientation during nest building in subjects with surgically removed eyes.

## Results and Discussion

In the first series of experiments performed in 2006, we tested a cohort (n = 6) of mole-rats before and after enucleation with the nest building assay in the ambient magnetic field. Before enucleation, the animals displayed a strong preference for the south-eastern sector of the arena (Figure 1 – figure supplement 1A; mean direction = 124° ± 53° (95% confidence interval), mean vector length r = 0.549, n = 6, Hotelling’s test: F = 15.610, p = 0.013), in line with published magnetic preferences in Ansell’s mole-rats (see Oliveriusová et al., 2012). In contrast, the distribution of nests built after enucleation was random (Figure 1 – figure supplement 1B; r = 0.140, n = 6, Hotelling’s test: F = 0.523, p = 0.628). These results suggested impairment of the magnetic sense, but did not allow us to exclude topographic factors or series effects.

**Figure 1:**
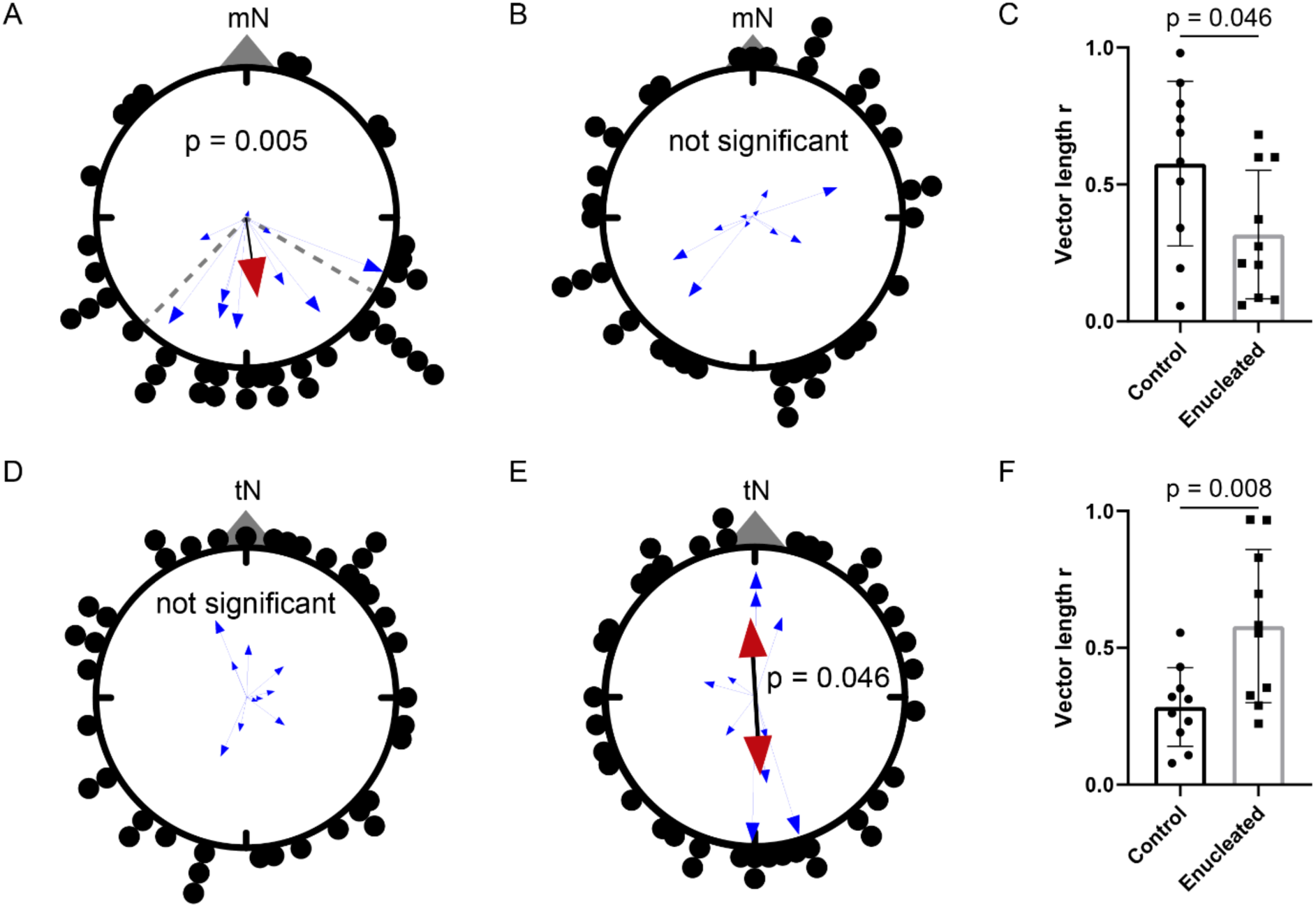
Enucleation results in a loss of magnetic directional preferences in the Ansell’s mole-rat. Nest distribution with respect to magnetic North (mN) of (A) controls and (B) enucleated Ansell’s mole-rats. Control animals (n = 10) exhibited a significant preference to build nests in the magnetic south-east, while the nests of enucleated animals (n = 10) were distributed randomly. (C) The mean angular vectors (calculated for each animal separately with respect to magnetic North) are significantly longer for controls compared to enucleated molerats. (D) There was no directional preference with respect to topographic north (tN) in control animals. (E) Enucleated mole-rats showed a weak but significant preference to build nests along the topographic north-south axis. (F) The mean angular vectors (calculated for each animal separately with respect to topographic North) are significantly longer for enucleated mole-rats compared to controls. The small blue arrows represent the mean vectors of four nests built by individuals or pairs of Ansell’s mole-rats and the arrow lengths reflect the r-values, a measure of the concentration of the nests. The red arrows are the weighted mean vector calculated over the mean vectors of tested individuals/pairs when significant. 95% confidence intervals are indicated by dashed lines. The dots outside of the circles indicate the positions of all 40 nests of each experimental group. The p-values indicate the results from Hotelling’s tests (A, E) or two-tailed t-tests (C, F) performed on the mean vectors.

To address these questions, we conducted a second series of experiments in 2018-2019, in which we tested a new cohort of enucleated (n = 10) and control (n = 10) animals with a coil setup that allowed us to precisely control the magnetic field. Furthermore, to minimize unspecific effects of the surgery, the behavioral experiments were performed more than 1.5 years after enucleation. We ascertained whether the enucleated animals behaved normally by recording ethograms of the experimental and control subjects in their home enclosures. The frequency measures of six quantified behaviors revealed no significant differences between enucleated subjects and controls (Figure 1 – figure supplement 2; Two-way ANOVA: F (1, 72) = 9.242×10^−18^, p > 0.999). In the nest building assay, all animals were tested in four different magnetic field alignments to distinguish magnetic from topographic responses. Expectedly, control animals displayed a significant preference for the magnetic south-eastern sector of the arena (Figure 1A; mean direction = 172° ± 52° (95% confidence interval), mean vector length r = 0.441, n = 10, Hotelling’s test: F = 11.027, p = 0.005). In contrast, the magnetic distribution of nests built by enucleated mole-rats was indistinguishable from random (Figure 1B; r = 0.129, n = 10, Hotelling’s test: F = 1.206, p = 0.348). The enucleated animals showed weaker magnetic orientation as indicated by the significantly shorter mean vectors compared to the control group (Figure 1C; mean r enucleated: 0.317, mean r control: 0.576, n = 10, two-tailed t-test: t = 2.146, df = 18, p = 0.046). With respect to topographic north, the nest directions of the controls did not deviate from a random distribution (Figure 1D; r = 0.112, n = 10, F = 0.4, p = 0.683), but a weak significant preference to build nests along the topographic north-south axis was found in the enucleated group (Figure 1E; mean direction = 176°/356°, r = 0.429, n = 10, F = 4.625, p = 0.046). Further, the mean vectors with respect to topographic north were significantly longer in the enucleated animals (Figure 1F; mean r enucleated: 0.580, mean r control: 0.284, two-tailed t-test: t = 2.970, df = 18, p = 0.008). These findings demonstrate that magnetic cues guided nest building in control animals, whereas enucleated animals appeared unable to perceive the magnetic field, leading them to fall back on individual topographic preferences.

In summary, we found a loss of magnetic directional preferences in two independent series of nest building experiments, yet we did not detect other behavioral consequences of enucleation in mole-rats. All enucleated animals were fully immersed members of their respective family groups and many successfully bred and raised offspring. As we carefully controlled for the influence of other sensory stimuli, we conclude that removal of the eyes led to permanent impairment of the magnetic sense. Alternatively, removal of the eyes might have had a yet uncharacterized neurological effect which could explain the lacking magnetic preferences, a possibility that should be addressed with experiments on a neuronal rather than behavioral level. The effect of enucleation on magnetic orientation is comparable to lidocaine application onto the cornea, corroborating the conclusion by Wegner et al. (2006) that the anesthesia affected the eyes rather than adjacent tissues or the central nervous system. Thus, future screens for light-independent, probably magnetite-based magnetoreceptors in this species should focus on the minute eyes of the Ansell’s mole-rat which are small enough (∼ 2 mm in diameter) to be visualized entirely by techniques such as high-throughput electron microscopy (compare Titze & Genoud, 2016; Graham et al., 2019). There have been attempts to detect magnetite in various rodent tissues (Mather, 1985) but, to our knowledge, no detailed surveys for ocular magnetite have been conducted in any mammal species so far. In transmission electron micrographs, Cernuda-Cernuda et al. (2003) noticed electron-dense crystalloid bodies in photoreceptors of the Ansell’s mole-rat and speculated that these consist of magnetite. They did, however, not quantify the size or distribution of the structures, nor did they try to confirm that the crystalloid bodies do contain iron. Wegner et al. (2006) identified ferric aggregates in the cornea by Prussian blue staining of a single mole-rat eye, which they interpreted as magnetite crystals. This intriguing, but non-replicated finding should be interpreted with caution. First, Prussian blue staining is not specific for magnetite. For example, iron-rich cells in the avian upper beak, once hypothesized to be magnetoreceptors based on this method, were later identified as macrophages, immune cells that accumulate ferric iron (Treiber et al., 2012). Although macrophages are typically absent from the rodent cornea, they invade corneal tissue during inflammation (Maruyama et al., 2005). Alternatively, iron contamination from the laboratory environment could explain the finding of Wegner et al. (2006) and must always be considered a potential confounder in histological screens for magnetoreceptors (Edelman et al., 2015).

If magnetite receptors are located in the cornea, they would most likely be innervated by the ophthalmic branch of the trigeminal nerve, which densely permeates the corneal tissue (Müller et al., 2003). This would be consistent with the finding of magnetically induced neural activity in a part of the Ansell’s mole-rat’s superior colliculus that predominantly receives trigeminal input (Němec et al., 2001). The trigeminal nerve has further been demonstrated to be involved in the magnetic sense in birds (Heyers et al., 2010, Lefeldt et al., 2014), but the exact location and structure of avian trigeminal magnetoreceptors is equally unknown (Engels et al., 2018). The presence of similar trigeminal magnetoreception systems in birds and mammals appears conceivable, but ophthalmic nerve ablation experiments coupled with behavioral assays are needed to establish the role of trigeminal input for magnetoreception in mole-rats. Our study highlights the significance of the eye for mammalian magnetoreception, which will facilitate future research on its cellular basis. Ultimately, this will contribute to characterize the function and significance of this enigmatic sensory channel within the mammalian radiation.

**Table 1:** Basic information on study animals used in series 1 (2006) and series 2 (2018/2019). Data on age correspond to the time of testing. µ and r represent the direction and lengths of the angular mean vector, respectively.

| ID | sex | age (months) | $\mu$ (control) | $r$ (control) | $\mu$ (enucl.) | $r$ (enucl.) |
| --- | --- | --- | --- | --- | --- | --- |
| 0287 | F | 30 | 119° | 0.99 | 285° | 0.32 |
| 3854 | M | 30 |  |  |  |  |
| 7556 | F | 82 | 89° | 0.68 | 250° | 0.11 |
| 0919 | M | 60 |  |  |  |  |
| 2668 | F | 47 | 182° | 0.59 | 41° | 0.38 |
| 3900 | M | 60 |  |  |  |  |
| 2848 | F | 72 | 127° | 0.46 | 150° | 0.31 |
| 3369 | M | 32 |  |  |  |  |
| 9546 | F | 56 | 134° | 0.58 | 37° | 0.32 |
| 3919 | M | 59 |  |  |  |  |
| 8118 | F | 91 | 105° | 0.43 | 106° | 0.67 |
| 0466 | M | 106 |  |  |  |  |

Series 2 (2018/2019)
| ID | sex | age (months) | enucleated/ control | $\mu$ | $r$ |
| --- | --- | --- | --- | --- | --- |
| 2654 | F | 55 | both enucleated | 47° | 0.06 |
| 0328 | M | 55 |  |  |  |
| 6125 | F | 66 | both enucleated | 125° | 0.21 |
| 6752 | M | 39 |  |  |  |
| 5496 | F | 69 | both enucleated | 71 ° | 0.60 |
| 0861 | M | 54 |  |  |  |
| 1334 | F | 75 | both enucleated | 250° | 0.27 |
| 0772 | M | 78 |  |  |  |
| 0772 | F | 31 | both control | 111° | 0.98 |
| 5813 | M | 43 |  |  |  |
| 6767 | F | 26 | both control | 162° | 0.83 |
| 0816 | M | 51 |  |  |  |
| 4780 | M | 69 | both control | 196° | 0.58 |
| 0726 | M | 21 |  |  |  |
| 0724 | F | 102 | both control | 150° | 0.51 |
| 0717 | M | 23 |  |  |  |
| 4295 | F | 57 | enucleated | 241° | 0.60 |
| 2175 | M | 65 | enucleated | 30° | 0.21 |
| 9074 | F | 90 | control | 245° | 0.34 |
| 1641 | M | 58 | enucleated | 118° | 0.37 |
| 5683 | F | 44 | enucleated | 218° | 0.68 |
| 0528 | F | 151 | control | 216° | 0.87 |
| 3360 | M | 184 | enucleated | 270° | 0.08 |
| 6126 | M | 42 | enucleated | 214° | 0.09 |
| 2731 | M | 27 | control | 18° | 0.06 |
| 0737 | F | 25 | control | 185° | 0.74 |
| 5222 | M | 26 | control | 121° | 0.19 |
| 3493 | M | 24 | control | 195° | 0.69 |

## Materials and Methods

### Subjects

We studied 40 (series 1: 6 males, 6 females; series 2:16 males, 12 females) adult Ansell’s molerats (*Fukomys anselli*) from the breeding stock of the Department of General Zoology at the University of Duisburg-Essen. All subjects were born in captivity and genealogically derived from individuals captured near Lusaka, Zambia. The animals were tested individually or as pairs (see Table S1). In enucleated animals, the eyes were removed entirely, ablating both the ophthalmic branch of the trigeminal nerve as well as the optic nerve. The enucleation procedure is described in detail in Henning et al. (2018). The recovery period between surgery and testing was a minimum of three weeks and 1.5 years in the first and second series of experiments, respectively. All surgeries and experiments conformed to the relevant ethical standards and were approved by the animal welfare officer of the University of Duisburg-Essen and the Landesamt für Natur, Umwelt und Verbraucherschutz NRW, Germany (series 1: 50.05-230- 37/06; series 2: 84-02.04.2015.A387).

### Ethograms

We recorded the behavior of four *F. anselli* families from experimental series 2 in their home terraria for a minimum of 435 minutes (maximum 1,338 minutes) using a Sony digital video camera recorder (HDR-CX505 VE or HDR-CX550VE). The families consisted of two, three, four and eight Ansell’s mole-rats, respectively, and included one or two enucleated mole-rats each (n = 6 enucleated). Eight out of ten non-enucleated mole-rats served as controls in the analysis; the behavior of two juveniles (< 12 months) was not evaluated. The analysis of the videos was performed blindly with respect to the experimental group (the resolution of the videos was not sufficient to resolve the mole-rats’ minute eyes). We evaluated the behavior for each individual separately on a minute by minute basis. The behavior that lasted longest during the respective minute was noted; rare events (grooming, conspicuous sniffing, or social play) were noted even if they occurred only shortly during the respective minute interval. The time budgets of the following behaviors were determined for each animal: resting, feeding (including food transport), locomotion (including digging), sniffing, grooming (auto-and allogrooming), and social play (e.g. sparring with their teeth, play fighting). The fraction of each behavioral category per total amount of recording time was calculated. Statistical differences between the experimental groups were tested using a two-way ANOVA with experimental group and type of behavior as factors (GraphPad Prism, V. 8.4.1).

### Nest-building assay

#### Testing site and experimental set-up

##### Experimental series 1

Animals were tested on warm days (ambient temperature 20-30 °C) in the year 2006, in an outside location of the Essen University campus in an undisturbed local geomagnetic field, i.e. within a greenhouse made from plastic walls with an aluminum frame (51°27’50.3”N 7°00’18.9”E, 48.6 µT, 66° inclination). Mole-rat pairs were tested in an opaque plastic arena (80·cm diameter, 30·cm height). The arena was covered with a thin layer of peat; tissue paper stripes and carrot pieces were spread radially across the floor. A bucket was placed in the middle of the arena to prevent central nesting. Animals were manually introduced into the arena at random directional segments. During testing, the arena was covered with a light impervious lid to exclude possible visual orientation. Each trial lasted up to 60 minutes after which the exact nest position was recorded with reference to geographic north. Each mole-rat pair underwent four tests under control conditions and six tests after enucleation.

##### Experimental series 2

The experiments took place between December 2018 and April 2019 in Essen-Haarzopf, a rural area with a stable magnetic field (49 µT, 66° inclination) in the periphery of the city of Essen, Germany (51°25’09.6”N 6°56’50.9”E). Experiments were conducted in a custom-build windowless wooden hut constructed completely of non-magnetic materials. The interior was shielded from radiofrequencies by a Faraday cage. Within the Faraday cage, the hut contained a double-wrapped three axial four coils Merritt system (edge length: 3 m x 3 m x 3 m), which was spatially arranged in a way to allow rotations of the magnetic field without changing inclination or intensity. The coils were powered by a programmable multichannel power supply (HMP4040 (384 W), Rohde & Schwarz, Munich, Germany) and magnetic field intensities were measured with a 3-axial fluxgate magnetometer (SENSYS FGM3D/75, SENSYS, Bad Saarow, Germany). Nest building took place in either of two circular black PVC arenas (arena 1: 80 cm (diameter) x 22 cm (height); arena 2: 65 cm (diameter) x 38 cm (height)) placed within a square wooden box (edge length: 120 cm). The arena was randomly turned before an experiment started to allow a yellow mark to be placed outside the arena (concealed from the animal’s view) was facing either topographic north or topographic south. This procedure reduced the likelihood that the animals orient by means of non-magnetic cues within the arena to which the experimenter might have been oblivious. The wooden box could be entered via four windows (size: 100 cm length x 80 cm (height), one on each cardinal direction, which were closed during the experiments. The arena was fixed on a pedestal seated in a sandbox to avoid substrate vibrations from irritating the animals during the experiments.

During testing, an LED table lamp emitting monochromatic red light (Parathom R50 80.337 E14 Red 617 nm, 6W, Osram, Munich, Germany) illuminated the room outside of the wooden box to allow the experimenters to operate. Ansell’s mole-rats are not capable of perceiving red light (Peichl et al., 2004). Photon density (light level) within the wooden box under testing conditions was measured at < 0.001 µmol s^-1^ m^-2^ with a photometer (LI-COR LI-250 Light Meter, LI-COR Biosciences, Lincoln, Nebraska). The temperature of the hut was controlled by an air conditioning system (Model FTXS50, Daikin) and was measured before (mean: 26.37 °C; SD: ± 1.74) and after (mean: 23.53 °C; SD: ± 2.06) each experimental run. The air conditioner was deactivated during experiments to minimize directional acoustic cues and prevent disruption of the magnetic environment within the hut.

#### Experimental procedure and scoring

We used the established nest building assay to study magnetoreception in the Ansell’s mole-rat (e.g., Marhold et al., 1997a; Wegner et al., 2006). Animals were placed singly or in pairs inside the arena (see Table S1) and were given approximately 60 min (mean: 67.7 min; SD: ± 11.6) to build a nest from paper scraps, which were evenly distributed along the outer wall of the arena at the beginning of the test. Traditionally, the nest building assay with mole-rats is performed with pairs, but to increase the number of statistically independent data points, we switched to testing individuals during the experiments. No animals were tested in both pairs and individually. Pairs and individual mole-rats were tested until four nests could be evaluated from each (see below). To evoke exploration of the arena, eight carrot slices were placed at the arena wall, aligned with the cardinal directions of the magnetic field. A white PVC cylinder (diameter 15 cm; height 20 cm) was placed in the middle of the arena to prevent nest construction in the center. Subjects were placed into the arena from either the (topographical) north, south or east door of the box following a randomized sequence. After placing the animal into the arena, the experimenter carefully closed the doors and immediately left the hut. After each test, the arena was carefully cleaned with mild detergent. To be able to distinguish topographic biases from magnetic orientation, all subjects had to complete four tests with different magnetic alignments (but unchanged topography). The animals were returned to their home enclosure for at least 24h between subsequent tests. During tests under simulated magnetic north conditions, antiparallel current was run through the Merritt coil system, so that possible side effects of the operational coils (noise, vibrations, heat, electric fields) were identical for all sessions. Thus, the other conditions differed only in the direction of magnetic north that was shifted by 90°, 180° or 270°, respectively (magnetic field intensities: north field: 47.76 µT, south field: 48.24 µT, east field: 48.12 µT, west field: 47.90 µT). To avoid order effects, the sequence of magnetic conditions was randomized for each animal. Four completed sessions were collected for each subject and pair, resulting in a total of 80 analyzed nest positions (40 enucleated, 40 controls).

A nest was considered valid for analysis when paper scraps were used to construct at least one clearly demarcated nest. In cases of multiple nest building, the vector of the largest nest was considered valid. Sessions were discarded when nests were built closer to the central cylinder then to the outer wall of the arena or when no nest was built. In case no unequivocal nest was constructed (n = 10 for controls; n = 8 for enucleated subjects), animals repeated trials under the respective magnetic condition until a valid nest was built. The nest positions were documented photographically.

### Data analysis

Nest orientations were scored (to the closest 5°) from photos by an experimenter blind to the magnetic condition and experimental group. Standard circular statistics were employed to analyze the distributions of the nest positions with respect to both magnetic and topographic orientation (Batschelet, 1981). All calculations were performed with Oriana 4.0 (Kovach Computing, Pentraeth, United Kingdom). Mean vectors of the four trials of each individual/pair were calculated with respect to magnetic or topographic north via vector addition. Second order statistics (Hotelling’s test) using mean vectors and lengths of the respective mean vectors were utilized to detect significant deviations from a random distribution, with α = 0.05. Two-tailed t-tests were used to test for differences in the length of the mean vectors of both experimental groups (GraphPad Prism V. 8.4.1).

## Acknowledgements

We thank Katharina Schröer for evaluating video recordings (behavioral analysis) and Georgina Fenton for stylistic comments on the manuscript. KRC was supported by a Ph.D. fellowship of the German National Academic Foundation (Studienstiftung des deutschen Volkes). This project was partly funded by the grant “EVA4.0”, No. CZ.02.1.01/0.0/0.0/16_019/0000803 financed by OP RDE of the European Union and the Ministry of Education, Youth and Sport of the Czech Rep (to SB).

