## Supplementary material for "Eyes are essential for magnetoreception in a mammal": Figure supplements

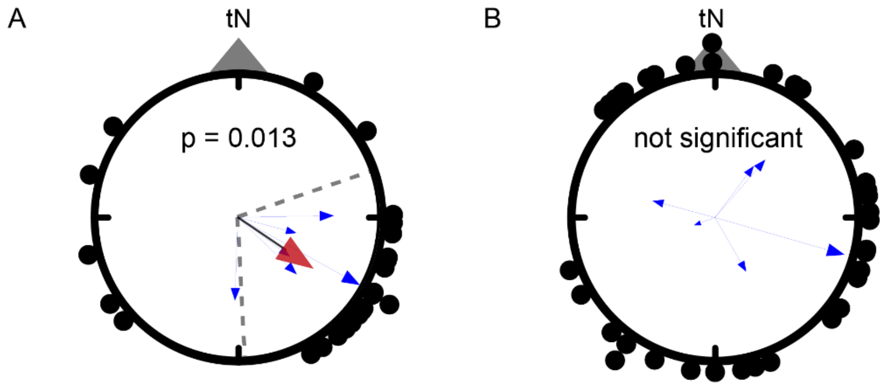


**Figure 1 – figure supplement 1: Pilot experiments showing a loss of directional preferences in the Ansell's mole-rat after enucleation**. Nest distribution of six pairs of Ansell's mole-rats (A) before (control) and (B) after surgery (enucleation). Control trials (n = 4 per pair) exhibited a significant preference to build nests in the magnetic south-east, while the nests built by the same pairs after surgery (n = 6 per pair) were distributed randomly. The small blue arrows represent the mean vectors of each pair of Ansell's mole-rats and the arrow lengths reflect the r-values, a measure of the concentration of the nests. The red arrows are the weighted mean vector calculated over the mean vectors of tested pairs when significant. 95% confidence intervals are indicated by dashed lines. The dots outside of the circles indicate the positions of all nests of each experimental group (controls: 24, enucleated: 36). The p-values indicate the results from Hotelling's tests performed on the mean vectors.

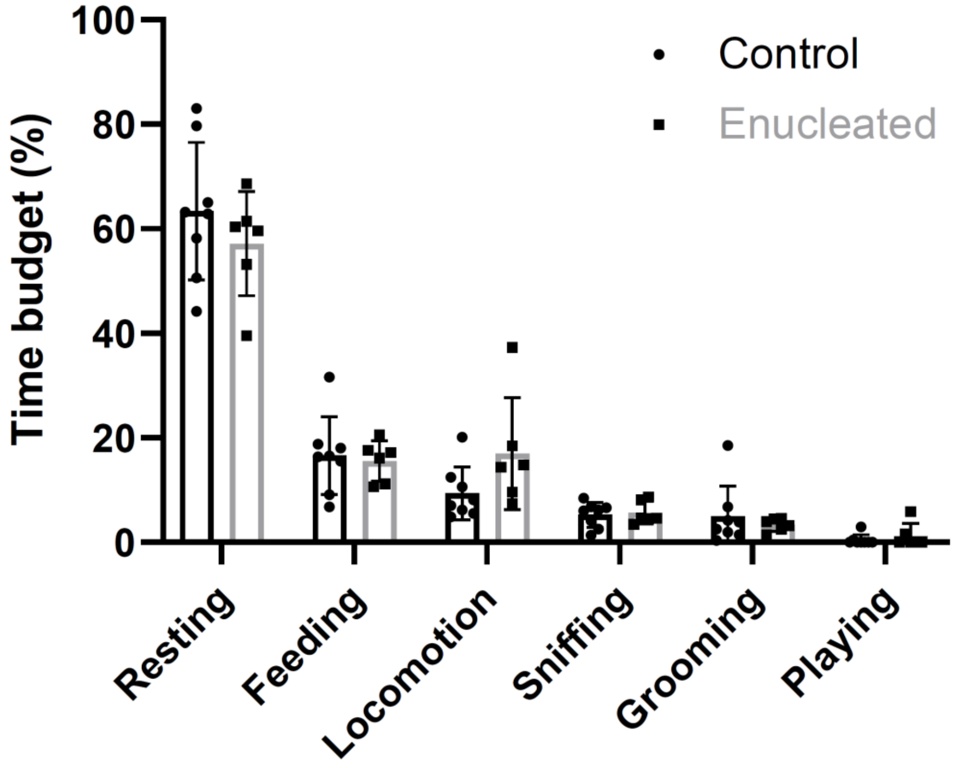


**Figure 1 – figure supplement 2: Enucleation does not affect the general behavior of Ansell's mole-rats.** Mean time budgets of six behaviors of enucleated and control Ansell's mole-rats observed in their home enclosures. All behaviors are expressed as the percentage of time spent per observation period. There were no significant differences between enucleated subjects and controls (Two-way ANOVA: F (1, 72) = 9.242x10 18, p > 0.999). n = 6 (enucleated) and n = 8 (controls), Error bars: standard deviation.
